# Gene dosage effect and reduced core fucosylation in B lymphocytes of *FUT8* heterozygous mice

**DOI:** 10.64898/2026.09.28.750819

**Authors:** Yue Ma, Jingxia Wang, Jingqing Liu, Shengyan Zhu, Nian Wang, Zhongxin Li, Dapeng Zhou

**Affiliations:** Department of Immunology and Pathogen Biology, Key Laboratory of Pathogen and Host Interactions, Ministry of Education, Tongji University School of Medicine, Shanghai, 200331, China; Autobio Diagnostics Co., Ltd., Zhengzhou 450016, China

**Keywords:** core fucosylation, FUT8, conditional knockout, genotyping, haploinsufficiency

## Abstract

Core fucosylation (α1,6-fucosylation) of N-glycans, catalyzed solely by fucosyltransferase 8 (FUT8), plays important roles in development, immunity, and antibody effector functions. Here we report the generation of *Fut8* floxed mice using a CRISPR/Cas9-based Extreme Genome Editing (EGE) system, in which loxP sites flank exon 8 of the *Fut8* gene and a LoxP-STOP-LoxP-RFP cassette was knocked in downstream of the promoter, rendering the targeted allele functionally null before Cre-mediated recombination. Crossing floxed mice with CMV-Cre transgenic mice yielded complete knockout mice carrying the Cre-recombined allele. A PCR-based genotyping method was established that unambiguously distinguishes the wild-type, heterozygous, and homozygous genotypes (+/+, Δ/+, and Δ/Δ). Intercrossing of heterozygous mice revealed that homozygous knockout mice die during the early postnatal period, consistent with previous reports, whereas heterozygous mice are viable. Flow cytometric analysis of AAL binding to splenic B cells and AAL lectin blotting both demonstrated markedly reduced core fucosylation in heterozygous mice (60.0 ± 6.8% and approximately 48% of wild-type levels, respectively). These results show that disruption of a single *Fut8* allele substantially reduces core fucosylation, and the mouse model described here provides a practical tool for functional studies of core fucosylation.

## Introduction

Core fucosylation (α1,6-fucosylation) is an important modification of N-glycans. FUT8 (α1,6-fucosyltransferase) is the sole enzyme in mammals that catalyzes this reaction, transferring fucose from GDP-fucose to the innermost N-acetylglucosamine (GlcNAc) of N-glycan chains via an α1,6-linkage in the Golgi apparatus [1]. Core fucosylation plays important roles in growth factor receptor signaling, growth and development, and immune regulation [2, 3, 10]. In addition, the absence of core fucose on the Fc region of IgG antibodies markedly enhances binding to FcγRIII and antibody-dependent cellular cytotoxicity (ADCC) [4]; therefore, inhibiting FUT8 is an effective approach to reduce antibody fucosylation. Since 2004, various strategies have been developed in cell engineering to reduce the core fucosylation of recombinant antibodies, including the generation of FUT8-knockout CHO cell lines [5-7], co-expression of GnTIII [8], and small-molecule fucosylation inhibitors [9].

To elucidate the physiological and pathological functions of core fucosylation, *Fut8* knockout mice have been generated and extensively characterized [10]. Approximately 70% of *Fut8* homozygous knockout mice die within 3 days after birth, and the survivors exhibit severe growth retardation and emphysema-like changes, which have been attributed to dysregulated activation and signaling of TGF-β1 and EGF receptors lacking core fucose [2, 3, 11]. Because of the low survival rate of homozygous knockout mice and the difficulty of obtaining adult homozygotes, heterozygous mice represent a practical research subject. Notably, Gao et al. reported that *Fut8* heterozygous mice exhibit markedly decreased FUT8 activity in the lung and increased susceptibility to cigarette smoke-induced emphysema [12], indicating that targeting a single allele is sufficient to impair core fucosylation. Nevertheless, systematic evaluation of core fucosylation levels in heterozygous mice remains limited.

Traditionally, targeted gene editing is achieved by homologous recombination, which is inefficient and time-consuming [13]. The CRISPR/Cas9 system recognizes targets through a synthetic guide RNA (sgRNA) and requires only a protospacer adjacent motif (PAM, 5’-NGG-3’) downstream of the target site; it is efficient and convenient and has been widely applied to the generation of knockout mice [14, 15].

In this study, we generated *Fut8* floxed mice using a CRISPR/Cas9-based Extreme Genome Editing (EGE) system: loxP sites were inserted into the introns flanking exon 8 of the mouse *Fut8* gene, and a LoxP-STOP-LoxP-RFP reporter cassette was knocked in downstream of the promoter. Owing to the insertion of the transcriptional STOP cassette, the targeted allele behaves as a loss-of-function allele before Cre-mediated recombination. Complete knockout mice were derived by crossing floxed mice with CMV-Cre transgenic mice, in which Cre-mediated recombination in the germline deleted the STOP cassette together with exon 8; the CMV-Cre transgene was subsequently segregated by breeding, and all mice used in subsequent experiments were Cre-negative. We established a PCR-based genotyping method that distinguishes wild-type, heterozygous, and homozygous genotypes (designated EGE-LYS-0112-A+/+, EGE-LYS-0112-AΔ/+, and EGE-LYS-0112-AΔ/Δ, respectively; hereafter +/+, Δ/+, and Δ/Δ, where Δ denotes the Cre-recombined knockout allele), and found by flow cytometry and Western blotting that core fucosylation levels in B lymphocytes of heterozygous mice were significantly lower than those in wild-type littermates. This model provides a useful tool for functional studies of core fucosylation and serves as a reference for the generation of knockout mouse models.

## Materials and Methods

### Mice

C57BL/6N mice were purchased from Biocytogen (Beijing, China) and housed under specific pathogen-free (SPF) conditions at the Experimental Animal Center of Tongji University, with free access to food and water. All animal experiments were approved by the Animal Ethics Committee of Tongji University (approval no. TJAA07421101). Age- and sex-matched wild-type (+/+) and heterozygous (Δ/+) littermates were used for experiments. All experimental mice were Cre-negative (Cre0), as confirmed by Cre genotyping.

### Generation of *Fut8* conditional knockout mice

The *Fut8* conditional knockout allele (designated EGE-LYS-0112-A) was generated using the Extreme Genome Editing (EGE) system, a CRISPR/Cas9-based genome editing technology with enhanced knock-in efficiency (Biocytogen, Beijing, China). Two sgRNAs were designed to target non-conserved regions within introns 7 and 8 of the mouse *Fut8* gene (chromosome 12, positive strand; NCBI ID: 53618). The targeting vector (12,085 bp) contained 5′ and 3′ homology arms of 1.4 kb and 1.3 kb, respectively, a loxP-flanked exon 8, and a LoxP-STOP-LoxP-RFP (LSL-RFP) reporter cassette inserted downstream of the promoter. Owing to the transcriptional STOP cassette, the targeted allele is transcriptionally silenced and thus behaves as a loss-of-function allele before Cre-mediated recombination; Cre-mediated excision removes the STOP cassette and exon 8, resulting in knockout of *Fut8* and RFP reporter expression. Cas9 mRNA, the two sgRNAs, and the targeting vector were microinjected into single-cell C57BL/6N zygotes, which were then transferred into pseudopregnant foster mothers. F0 founder mice were screened by PCR and confirmed by Southern blotting and Sanger sequencing. F1 mice carrying the targeted allele (fl/+) were generated by crossing founders with wild-type C57BL/6N mice, and homozygous floxed mice (fl/fl) were obtained by intercrossing F1 mice. To generate complete knockout mice, fl/fl mice were crossed with CMV-Cre transgenic mice (Biocytogen, Beijing, China), in which Cre recombinase is ubiquitously expressed; Cre-mediated recombination in the germline deleted the STOP cassette together with exon 8, and the resulting knockout allele (Δ) was transmitted through the germline. After the Cre transgene was segregated out by crossing to wild-type mice, heterozygous knockout mice (Δ/+) were intercrossed to obtain homozygous complete knockout mice (Δ/Δ) (Fig. S2). Sperm from homozygous floxed (fl/fl) male mice were cryopreserved for future tissue-specific conditional knockout applications.

### Genotyping

Genomic DNA was extracted from tail tips (21-day-old mice) or embryonic tissues using the TransDirect Animal Tissue PCR Kit (TransGen Biotech, Beijing, China) according to the manufacturer’s instructions (lysis at 55 °C for 30 min, followed by 95 °C for 3 min). Three primer pairs were used for genotyping (Table 1). PCR was performed in a single 10-μL triplex reaction containing 2.8 μL ddH2O, 5 μL 2× DirectMix, 0.6 μL EGE-LYS-0112-A-5′loxP-F (10 μM), 0.3 μL EGE-LYS-0112-A-3′loxP-R (10 μM), 0.3 μL EGE-LYS-0112-A-5′loxP-R (10 μM), and 1 μL DNA template. The cycling conditions were: 94 °C for 2 min; 35 cycles of 98 °C for 10 s, 62 °C for 30 s, and 68 °C for 1 min/kb; and a final extension at 68 °C for 10 min. PCR products were separated on 1.8–2% agarose gels containing 0.01% nucleic acid stain at 120 V for 30 min and visualized with a gel imaging system (CLINX). The expected band patterns were: wild-type (+/+), 362 bp; floxed heterozygous (fl/+), 362 bp and 456 bp; floxed homozygous (fl/fl), 456 bp; knockout heterozygous (Δ/+), 362 bp and 751 bp; knockout homozygous (Δ/Δ), 751 bp; the 751-bp product corresponded to the Cre-recombined knockout allele (Δ). Cre genotyping was performed in a separate PCR reaction using the Cre-F4/Cre-R4 primer pair (385-bp product) under the same cycling conditions.

**Table 1.**
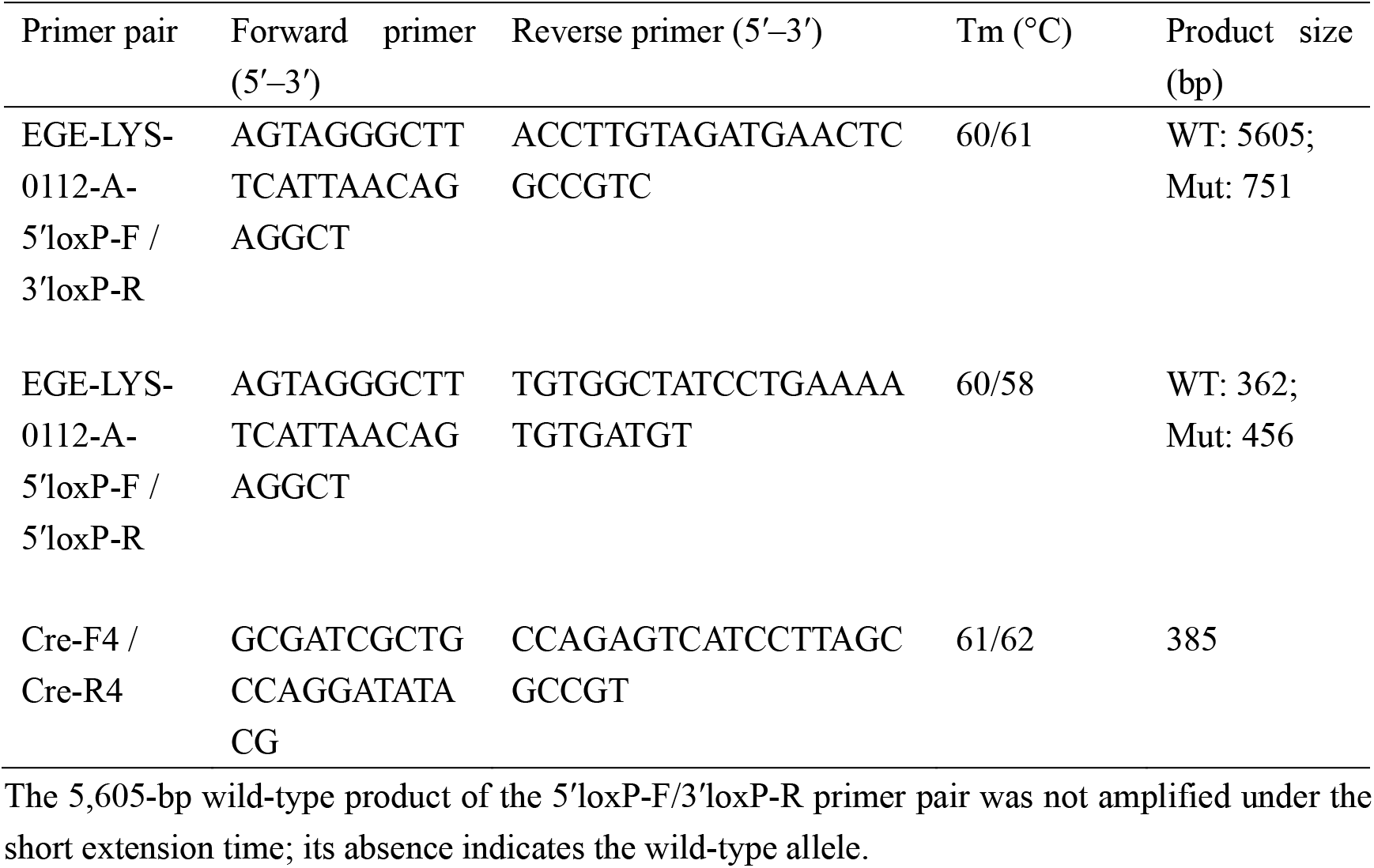
Primers used for genotyping.

### Embryo collection and genotyping

Heterozygous (Δ/+) mice were set up for timed mating, and embryos were collected at embryonic days E18–E20 after plug check. Yolk sacs or embryonic tissues were dissected for genomic DNA extraction, and embryos were genotyped as described above (8 embryos in total). To determine the fate of homozygous knockout mice, newborn pups were monitored and dead pups were collected for genotyping; 22 pups from two litters were sampled at 15–20 days of age, of which 3 were Δ/Δ and were already dead at the time of collection.

### Flow cytometry

Splenocytes were prepared from 8-to 12-week-old wild-type (+/+) and heterozygous (Δ/+) littermates; all experimental mice were Cre-negative (Cre0). B cells were isolated using a mouse B cell isolation kit (cat# 19844; STEMCELL Technologies) according to the manufacturer’s instructions. Isolated B cells were incubated with biotinylated AAL (2 μg/mL; cat# B-1395-1; Vector Laboratories) at 4 °C for 30 min in the dark, washed, and then incubated with APC-conjugated streptavidin (1 μg/mL; cat# 17-4317-82; Invitrogen eBioscience) and FITC-conjugated anti-mouse CD45R/B220 (2 μg/mL; cat# 553088; BD Pharmingen) at 4 °C for 30 min. After washing, samples were acquired on a BD LSRFortessa flow cytometer (BD Biosciences) and analyzed with FlowJo software (v10). The median fluorescence intensity (MFI) of AAL-APC staining was compared between genotypes within the B220+ B cell population. For the AAL-negative B cell analysis (Fig. S1), splenocyte suspensions were stained as described above, without prior magnetic B cell isolation. Secondary antibody-only and unstained samples served as controls.

### AAL lectin blotting

Total protein was extracted from splenic B cells of 8-to 12-week-old +/+ and Δ/+ mice (isolated by magnetic cell sorting as described above) using RIPA lysis buffer containing protease inhibitors, and protein concentrations were determined by a BCA assay. Equal amounts of protein (2 μg per lane) were separated by 10% SDS-PAGE and transferred onto PVDF membranes. Membranes were blocked with 2% BSA at room temperature for 2 h, incubated with biotinylated AAL (1:10,000 dilution in 2% BSA; cat# B-1395-1; Vector Laboratories) overnight at 4 °C, and then incubated with HRP-conjugated streptavidin (1:10,000 dilution in 2% BSA) at room temperature for 1 h. Signals were visualized by enhanced chemiluminescence, and band intensities were quantified with ImageJ and normalized to GAPDH (loading control).

### Statistical analysis

Data are presented as mean ± SEM. Statistical analyses were performed using GraphPad Prism 8.0. Two-group comparisons were made by unpaired two-tailed Student’s t tests, normalized flow cytometry percentages were compared with 100% by one-sample t test, and genotype distributions were analyzed by the chi-square test. P < 0.05 was considered statistically significant.

## Results

### Generation of *Fut8* conditional knockout mice

To investigate the physiological roles of core fucosylation in vivo, we generated *Fut8* floxed mice using the CRISPR/Cas9-based EGE system. loxP sites were inserted into the introns flanking exon 8 of the mouse *Fut8* gene, and a LoxP-STOP-LoxP-RFP reporter cassette was knocked in downstream of the promoter (Fig. 1A). Because the transcriptional STOP cassette silences the targeted allele, EGE-LYS-0112-A behaves as a loss-of-function allele prior to Cre-mediated recombination; Cre-mediated excision removes the STOP cassette and exon 8, resulting in *Fut8* knockout and RFP reporter expression. F0 founders were confirmed by PCR, Southern blotting, and Sanger sequencing. F1 mice carrying the floxed allele (fl/+) were obtained by crossing F0 founders with wild-type C57BL/6N mice. To achieve complete knockout, homozygous floxed (fl/fl) mice were crossed with CMV-Cre transgenic mice, in which Cre recombinase is ubiquitously expressed; Cre-mediated recombination in the germline deleted the STOP cassette together with exon 8, and the knockout allele (Δ) was transmitted through the germline. After the CMV-Cre transgene was segregated out by crossing to wild-type mice, heterozygous knockout mice (Δ/+) were intercrossed to obtain homozygous complete knockout mice (Δ/Δ) (Fig. S2). All mice used in subsequent analyses were Cre-negative (Cre0), as confirmed by genotyping. In parallel, the floxed line was preserved (sperm from fl/fl males cryopreserved) for future tissue-specific conditional knockout applications.

**Figure 1.**
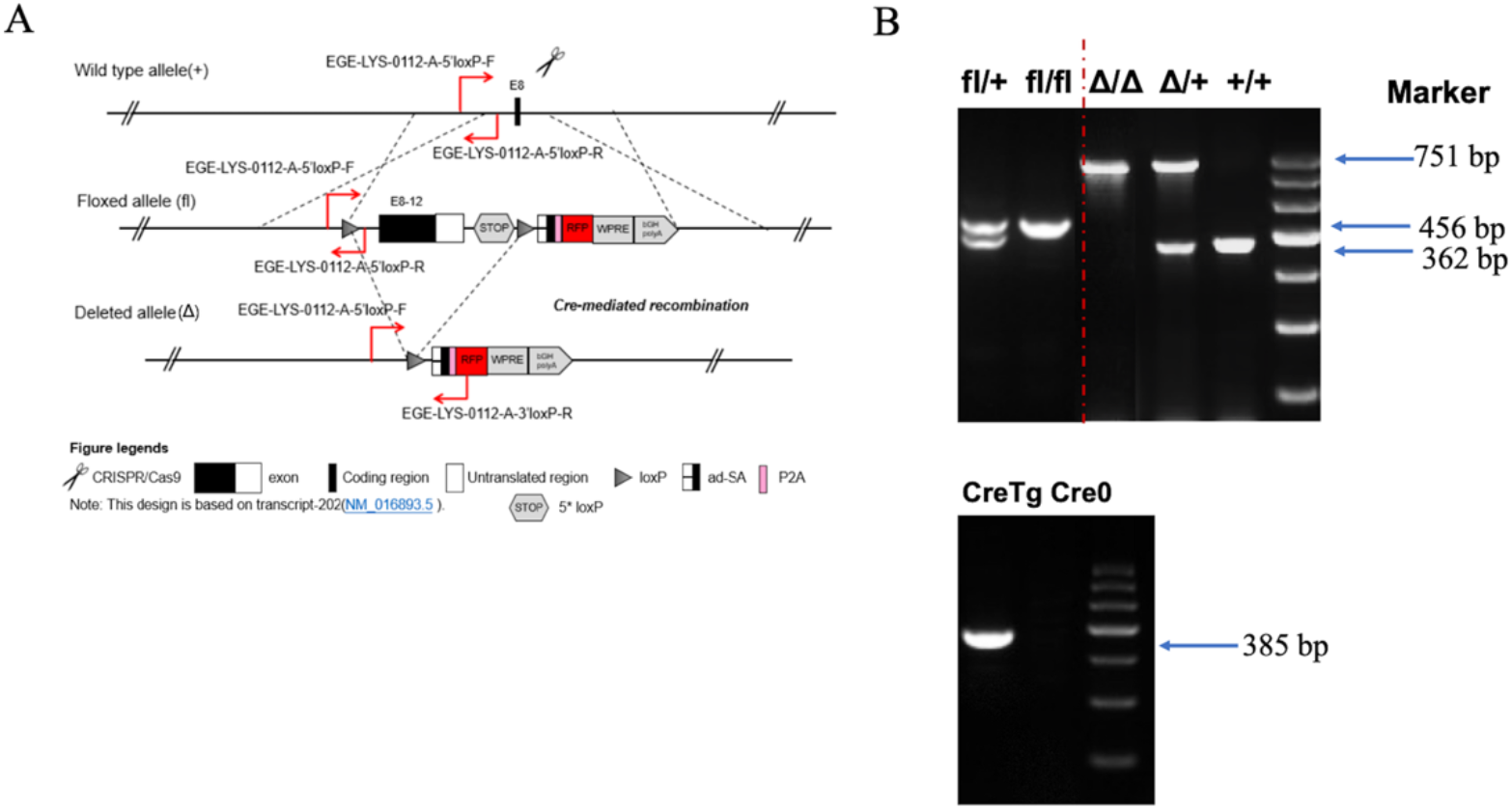
Generation and genotyping of *Fut8* floxed and complete knockout mice. (A) Targeting strategy for the EGE-LYS-0112-A allele: loxP sites were inserted into introns 7 and 8 flanking exon 8, and a LoxP-STOP-LoxP-RFP cassette was knocked in downstream of the promoter. (B) Representative genotyping PCR. All genotype symbols refer to the Fut8 locus: +/+, wild-type; fl/+ and fl/fl, heterozygous and homozygous floxed mice (exon 8 flanked by loxP but not deleted); Δ/+, heterozygous complete knockout; Δ/Δ, homozygous complete knockout. Δ denotes the Cre-recombined knockout allele, in which exon 8 (together with the STOP cassette) has been deleted. All mice used in subsequent experiments were Cre-negative (Cre0): the CMV-Cre transgene had been segregated by breeding after germline recombination. Top: all lanes were amplified in a single triplex PCR reaction containing three primers (5′loxP-F, 5′loxP-R, and 3 ′ loxP-R), whose binding sites are indicated in (A); each lane shows the products corresponding to the alleles carried by that mouse. Lanes (left to right): fl/+ (362 and 456 bp), fl/fl (456 bp), Δ/Δ (751 bp), Δ/+ (362 and 751 bp), +/+ (362 bp), and DNA size marker. The dashed line separates the two mouse lines: lanes to its left show mice of the floxed line (fl/+ and fl/fl), and lanes to its right show mice of the complete knockout line (Δ/Δ and Δ/+) together with a +/+ control. The 362-bp product corresponds to the wild-type allele, the 456-bp product to the non-deleted floxed allele, and the 751-bp product to the Cre-recombined knockout allele (Δ). Bottom: PCR detection of the CMV-Cre transgene in a separate reaction, illustrating the presence (CreTg) and absence (Cre0) of the transgene; lanes (left to right): CreTg (385 bp), Cre0, and DNA size marker.

### Establishment of a PCR-based genotyping method

To genotype mice carrying the targeted allele, we established a PCR-based method in which the flox- and knockout-detection primers were combined in a single triplex reaction (Table 1). The wild-type allele yields a 362-bp product, the knockout allele a 751-bp product, and the non-deleted floxed allele a 456-bp product; the Cre transgene is detected by a 385-bp product (Fig. 1B, bottom). This strategy unambiguously distinguishes +/+ (362 bp), fl/+ (362 and 456 bp), fl/fl (456 bp), Δ/+ (362 and 751 bp), and Δ/Δ (751 bp) mice (Fig. 1B).

### Homozygous *Fut8* knockout mice die during the early postnatal period

Heterozygous mice were intercrossed, and timed-pregnant females were dissected at E18–E20. Among 8 embryos, 4 were Δ/+ and 4 were Δ/Δ (χ^2^ = 4.0, df = 2, P = 0.14 versus the expected 1:2:1 ratio, Table 2, Fig. S3A), consistent with embryonic viability of Fut8-null mice through E18–E20. By contrast, among 22 pups genotyped at 15–20 days of age from two litters, 10 were +/+, 9 were Δ/+, and only 3 were Δ/Δ (13.6% versus 25% expected; χ^2^ = 5.18, df = 2, P = 0.075) (Table 2), and all three Δ/Δ pups were already dead at the time of collection. Early postnatal death was observed in the Δ/Δ animals (Table 2, Fig. S3B), as expected for complete Fut8 knockout. These results indicate that homozygous *Fut8* knockout mice die during the early postnatal period, consistent with the neonatal lethality reported for *Fut8*-null mice [2], whereas heterozygous mice survive to adulthood.

**Table 2.** Genotype distribution of offspring from heterozygous (Δ/+) intercrosses.

| Offspring | Total | +/+ | $\Delta/+$ | $\Delta/\Delta$ | $\chi^2$ (df = 2) | P |
| --- | --- | --- | --- | --- | --- | --- |
| Embryos (E18–E20) | 8 | 0 | 4 | 4 | 4.0 | 0.14 |
| Pups (15–20 days) | 22 | 10 | 9 | 3 | 5.18 | 0.075 |

$\chi^2$ values were calculated against the expected 1:2:1 Mendelian ratio. All three $\Delta/\Delta$ pups were already dead at the time of collection. All offspring were Cre-negative (Cre0).

### Reduced core fucosylation in heterozygous mice

To assess whether disruption of a single *Fut8* allele affects core fucosylation, we compared α1,6-core fucosylation levels between Δ/+ mice and +/+ littermates using flow cytometry and AAL lectin blotting. In flow cytometry, AAL binding to splenic B cells from Δ/+ mice was 60.0 ± 6.8% of that in +/+ controls (mean ± SEM, n = 3 independent experiments; one-sample t test versus 100%, P = 0.028) (Fig. 2A; Table 3). In addition, a small AAL-negative B cell population was detected among splenocytes of Δ/+ mice (3.78%) but was minimal in +/+ controls (0.56%) (Fig. S1). Consistently, AAL lectin blotting of splenic B cell lysates showed markedly reduced signal across the molecular weight range in Δ/+ samples; whole-lane densitometry normalized to GAPDH indicated that total AAL signal in Δ/+ samples was approximately half of that in +/+ controls in two independent experiments (Fig. 2B; Table 4). Together, these results demonstrate that core fucosylation is substantially reduced in heterozygous *Fut8* knockout mice, indicating a gene-dosage effect (haploinsufficiency).

**Table 3.** AAL binding to splenic B cells from Δ/+ mice relative to +/+ controls.

| Experiment | +/+ MFI | $\Delta/+$ MFI | $\Delta/+$ (% of +/+) |
| --- | --- | --- | --- |
| 1 | 77,195 | 38,275 | 49.6 |
| 2 | 33,037 | 19,031 | 57.6 |
| 3 | 26,021 | 18,921 | 72.7 |
| Mean ± SEM | – | – | 60.0 ± 6.8 |

MFI, mean fluorescence intensity. Each experiment used one +/+ and one $\Delta/+$ mouse; all experimental mice were Cre-negative (Cre0). The normalized percentages were compared with 100% by one-sample t test ( $t = 5.9$ , $df = 2$ , $P = 0.028$ ).

**Table 4.** AAL lectin blot densitometry of splenic B cell lysates from Δ/+ mice relative to +/+ controls.

| Experiment | AAL ($\Delta/+$ / $+/+$) | GAPDH ($+/+$ / $\Delta/+$) | Normalized $\Delta/+$ (% of $+/+$) |
| --- | --- | --- | --- |
| 1 | 0.27 | 1.90 | 51 |
| 2 | 0.49 | 0.90 | 44 |
| Mean | — | — | 48 |

Whole-lane densitometry of AAL signal, normalized to GAPDH (ImageJ). Normalized values = AAL ratio × GAPDH ratio. One $+/+$ and one $\Delta/+$ mouse were analyzed per experiment; all experimental mice were Cre-negative (Cre0).

**Figure 2.**
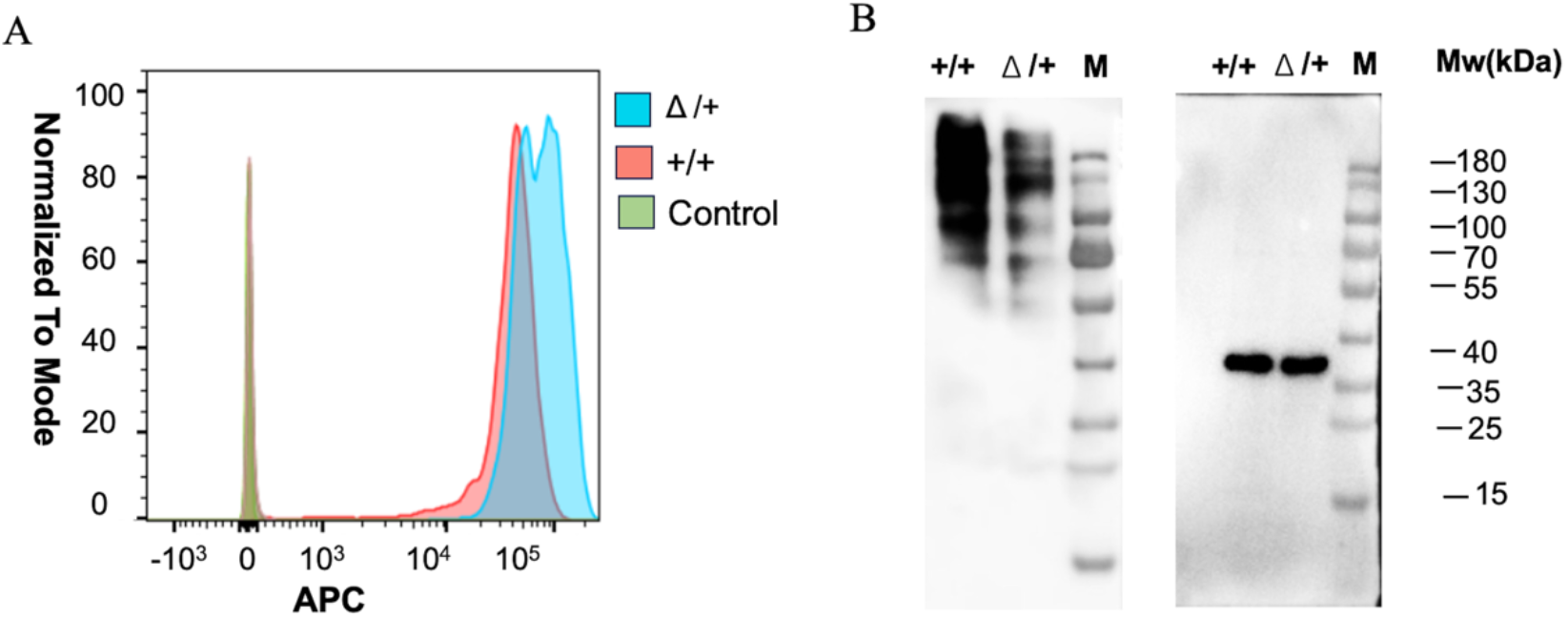
Reduced core fucosylation in heterozygous *Fut8* knockout mice. (A) Flow cytometric analysis of AAL binding to B220+ splenic B cells from +/+ and Δ/+ mice; representative of three independent experiments; quantification is shown in Table 3. (B) AAL lectin blot of splenic B cell lysates (2 μg per lane) with GAPDH as a loading control (left: AAL; right: GAPDH). Lanes in each blot: +/+ (left), Δ/+ (middle), and DNA size marker (right). Representative of two independent experiments; quantification is shown in Table 4.

## Discussion

The reduced core fucosylation in heterozygous mice is in line with a previous report that heterozygous *Fut8* knockout mice exhibit markedly decreased FUT8 activity in the lung and increased susceptibility to cigarette smoke-induced emphysema [12]. Our results extend these findings by showing that heterozygous mice exhibit markedly reduced core fucosylation in splenic B cells (approximately half of wild-type levels, as measured by two independent methods), and that the reduction is consistent with the loss of one functional *Fut8* allele. The slightly greater reduction detected by lectin blotting than by flow cytometry likely reflects differences between total cellular and cell-surface glycoproteins.

Whereas the complete knockout mice in this study were derived with the ubiquitously expressed CMV-Cre, the cryopreserved floxed line additionally enables tissue-specific conditional knockout: the LSL-RFP cassette allows Cre-dependent RFP labeling of *Fut8*-deficient cells, and tissue-specific or inducible Cre lines can be used to inactivate *Fut8* in adult mice, circumventing the early postnatal lethality of germline homozygotes. A limitation of this study is the modest sample size for the embryo and pup collections and the limited number of independent experiments; nevertheless, the consistency among flow cytometry, lectin blotting, and the lethality data supports the robustness of the conclusions. In summary, we generated and validated *Fut8* floxed and complete knockout mouse models and established a reliable genotyping strategy, providing a useful tool for investigating the functions of core fucosylation in vivo.

## Data availability

All data supporting the findings of this study are available within the manuscript. Mouse lines and other resources are available from the corresponding author upon reasonable request.

## Author contributions

Y.M. performed experiments, analyzed data, and wrote the manuscript. J.W., J.L., S.Z., and N.W. performed experiments. Z.L. provided resources. D.Z. conceived and supervised the study. All authors read and approved the final manuscript.

## Figure Legends

**Figure S1.**
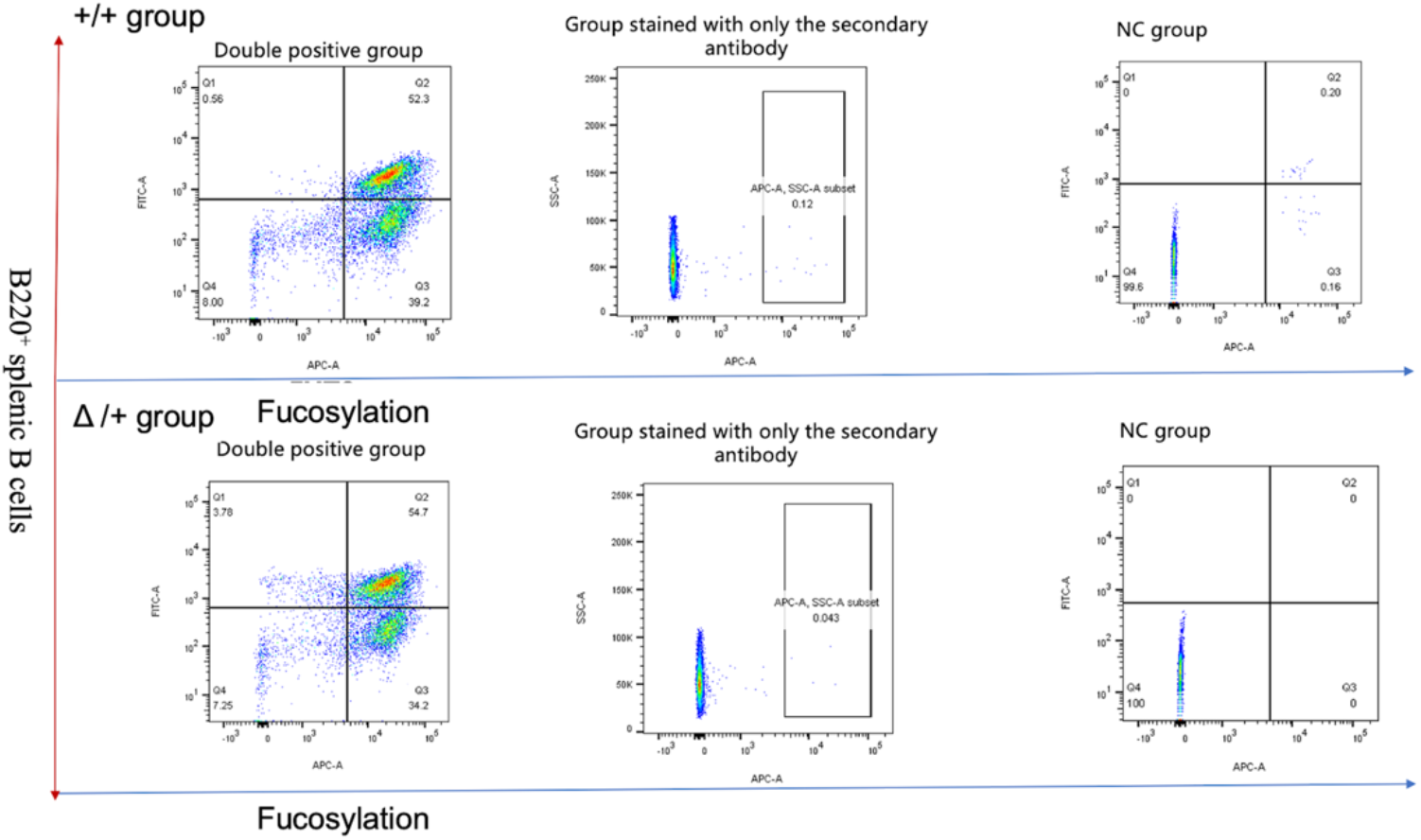
Flow cytometric analysis of AAL-negative B cells among splenocytes of +/+ and Δ/+ mice. Splenocytes were stained with FITC-conjugated anti-B220 and biotinylated AAL followed by streptavidin–APC. Dot plots are shown for +/+ mice (top row) and Δ/+ mice (bottom row), in identical order from left to right: full staining, secondary antibody only (streptavidin–APC without biotinylated AAL), and unstained negative control. The AAL-negative B cell population accounted for 0.56% in +/+ mice and 3.78% in Δ/+ mice, indicating the emergence of a small B cell subpopulation lacking detectable core fucosylation in heterozygous mice.

**Figure S2.**
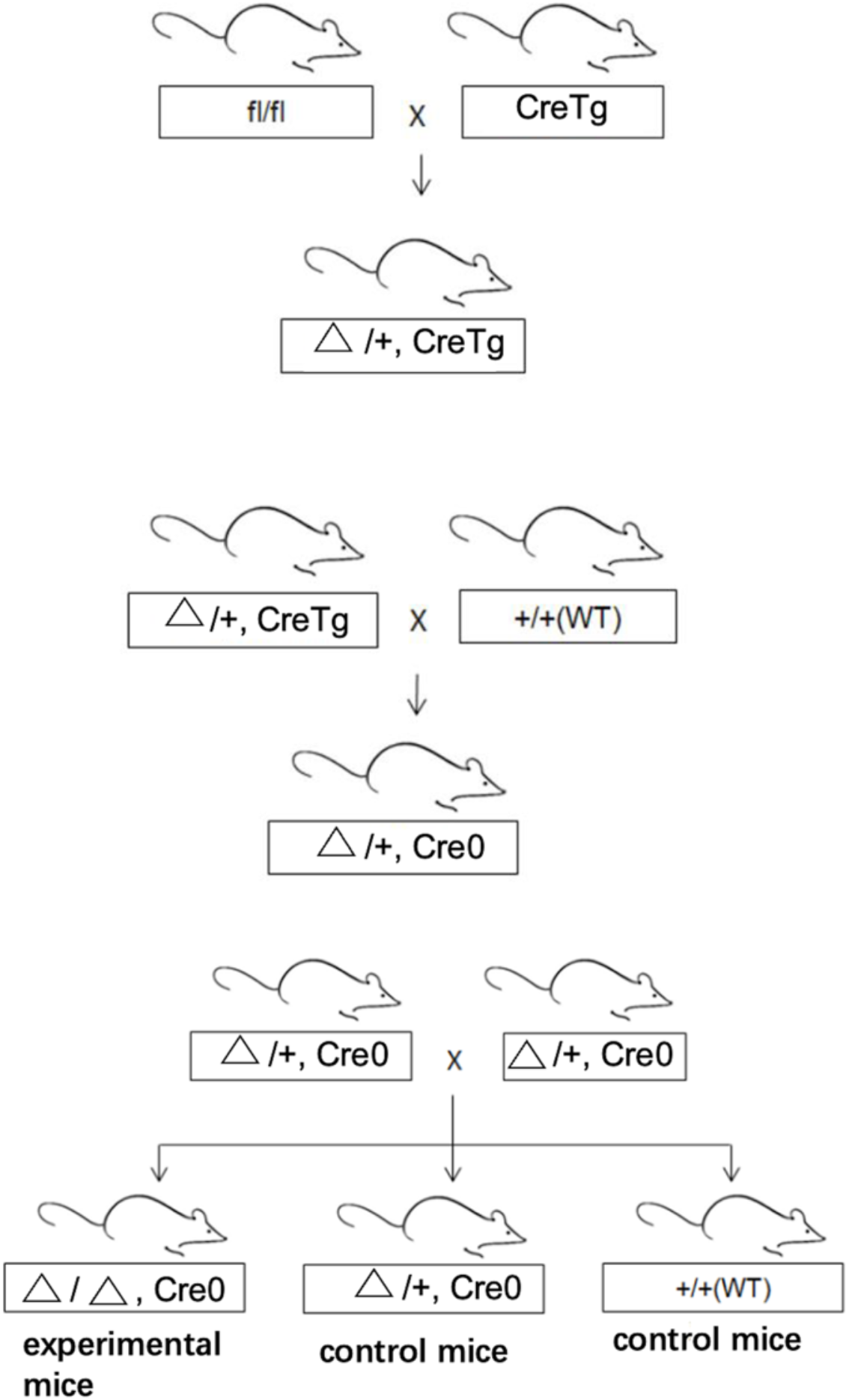
Breeding scheme for the generation of complete *Fut8* knockout mice. Homozygous floxed mice (fl/fl) were crossed with CMV-Cre transgenic mice (CreTg), and Cre-mediated recombination in the germline deleted the STOP cassette together with exon 8, generating the knockout allele (Δ). The resulting Δ/+;CreTg mice were crossed with wild-type mice to segregate out the Cre transgene, and Cre-negative (Cre0) Δ/+ mice were selected and intercrossed to obtain Δ/Δ mice.

**Figure S3.**
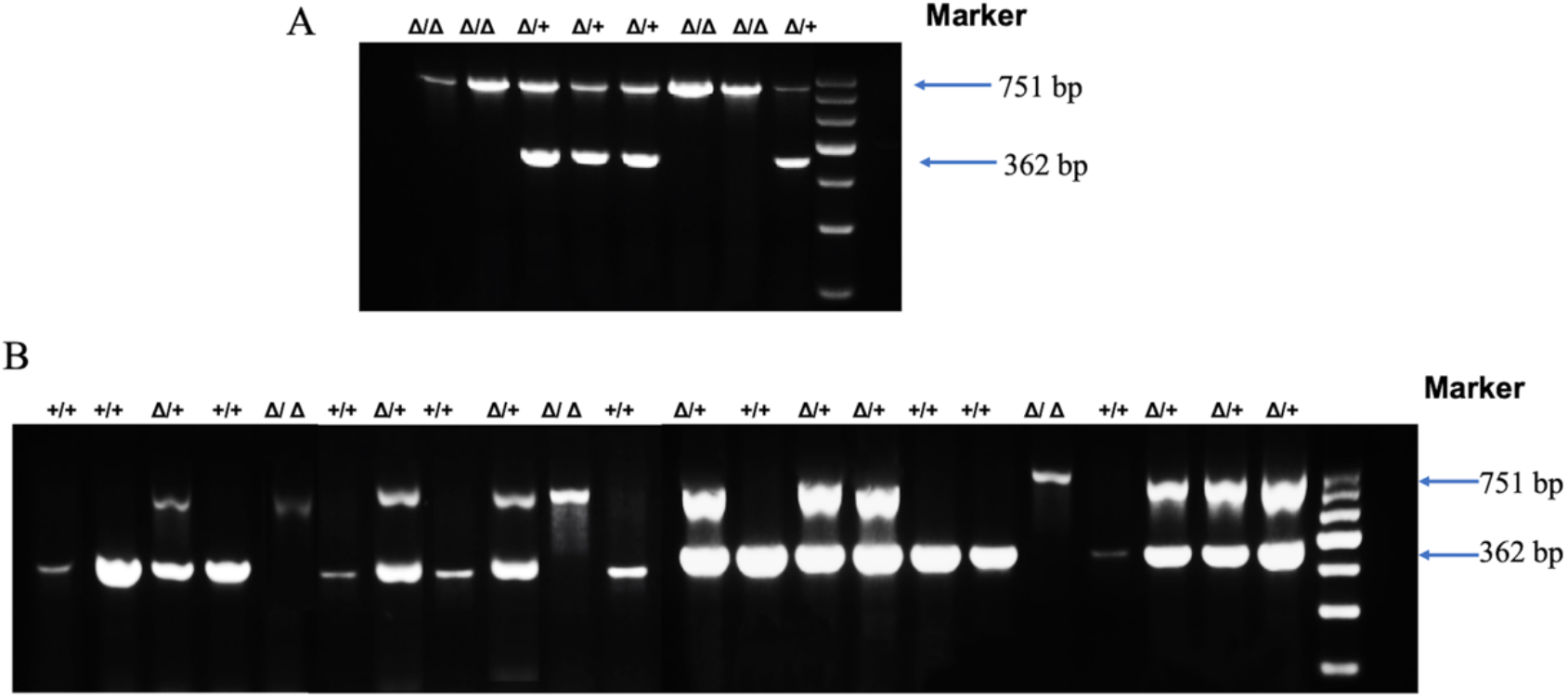
Genotyping of intercross progeny by triplex PCR. (A) Representative genotyping gel of E18–E20 embryos from heterozygous intercrosses. The wild-type allele yields a 362-bp product and the knockout allele a 751-bp product; among the 8 embryos collected, 4 were Δ/+ (362 and 751 bp) and 4 were Δ/Δ (751 bp) (Table 2). (B) Representative genotyping gel of pups collected at 15–20 days of age from two litters. Ten +/+, 9 Δ/+, and 3 Δ/Δ pups were identified (Table 2); the three Δ/Δ pups were already dead at the time of tail-tip collection.

